# Allelic diversity uncovers protein domains contributing to the emergence of antimicrobial resistance

**DOI:** 10.1101/2022.10.24.513455

**Authors:** Trudy-Ann Grant, Mario López-Pérez, Salvador Almagro-Moreno

## Abstract

Antimicrobial resistance (AMR) remains a major threat to global health. To date, tractable approaches that decipher how AMR emerges within a bacterial population remain limited. Here, we developed a framework that exploits genetic diversity from environmental bacterial populations to decode emergent phenotypes such as AMR. OmpU, is a porin that makes up to 60% of the outer membrane of *Vibrio cholerae*, the cholera pathogen. This porin is directly associated with the emergence of the bacterium and confers resistance to numerous host antimicrobials. In this study, we examined naturally occurring allelic variants of OmpU in environmental *V. cholerae* and established associations that connected genotypic variation with phenotypic outcome. We covered the landscape of gene variability and found that the porin forms two major phylogenetic clusters with striking genetic diversity. We generated 14 isogenic mutant strains, each encoding a unique *ompU* allele, and found that divergent genotypes lead to convergent antimicrobial resistance profiles. We identified and characterized functional domains in OmpU unique to variants conferring AMR-associated phenotypes. Specifically, we identified four conserved domains that are linked with resistance to bile and host-derived antimicrobial peptides. Mutant strains for these domains exhibit differential susceptibility patterns to these and other antimicrobials. Interestingly, a mutant strain in which we exchanged the four domains of the clinical allele for those of a sensitive strain exhibits a resistance profile closer to a porin deletion mutant. Finally, using phenotypic microarrays, we uncovered novel functions of OmpU and their connection with allelic variability. Our findings highlight the suitability of our approach towards dissecting the specific protein domains associated with the emergence of AMR and can be naturally extended to other bacterial pathogens and biological processes.

**AUTHOR SUMMARY:** Antimicrobial resistance (AMR) is one of the major threats to global health. To date, tractable approaches that decipher how AMR emerges within a bacterial population remain limited. Here we developed an approach that uses genetic diversity from environmental populations to decode emergent phenotypes such as AMR. Specifically, we examined naturally occurring allelic variants of an outer membrane porin, OmpU, in *Vibrio cholerae* and established associations between genotype and phenotype. Using this approach, we identified and characterized the functional domains in OmpU unique to variants conferring AMR-associated phenotypes. Our perspective towards disentangling the emergence of AMR can be naturally extended to other proteins and bacterial pathogens.

## INTRODUCTION

Antimicrobial resistance (AMR) is a complex phenomenon fueling the arms race to combat emerging and re-emerging pathogens. By 2050, it is estimated that 10 million individuals will die annually due to infections caused by pathogens that exhibit AMR [1]. Resistance to many currently used antimicrobials such as ciprofloxacin or ampicillin has emerged in up to 80% of *Klebsiella pneumoniae* and *Escherichia coli* strains, increasing the likelihood of treatment failures [2]. Current approaches to combat AMR include the development of antimicrobial cocktails and the discovery of novel or modified ones, however, the rapid adaptations to new therapeutics by pathogenic strains demands a deeper understanding of the molecular mechanisms and evolutionary paths associated with the emergence of this threat [3, 4]. Furthermore, given the complexity of this problem, approaches that recognize the relevance of the environment that pathogenic agents inhabit and closely align with the “One Health” paradigm are imperative to effectively address AMR [5-8].

Gram-negative bacteria pose one of the greatest challenges to tackle AMR due to their highly selective outer membrane embedded with proteins that hinder the effectiveness of antimicrobial compounds [9-11]. One group of membrane proteins commonly associated with AMR are outer membrane porins (OMPs). OMPs facilitate the translocation of molecules within the cell and are often essential to pathogenesis as regulators of this selective barrier [12-15]. OMPs modulate transport of small molecules, with selectivity based upon charge and size, allowing for the intake of essential nutrients, or preventing internalization of antimicrobials [16-20]. Furthermore, OMPs confer antimicrobial resistance through alteration in levels of porin expression, mutations in the porin constriction region, or expression of alternative porins [15, 18, 21-26]. To date, how these critical adaptations for virulence evolve remains poorly understood.

*Vibrio cholera*e, a comma-shaped Gram-negative bacterium, is the etiological agent of the severe diarrheal disease cholera. Interestingly, only a phylogenetically confined group within the species, the pandemic cholera group (PCG), can cause the disease in humans [27-29]. Recently, we determined that strains from PCG encode allelic variations in core genes that confer preadaptations for virulence [27]. These allelic variations of core genes, termed virulence adaptive polymorphisms (VAPs), naturally circulate in environmental populations of *V. cholerae* including non-toxigenic strains [27]. OmpU is a major porin in *V. cholerae* that can make up to 60% of its outer membrane and is associated with antimicrobial resistance and intestinal colonization, among other phenotypes [30-33]. Recently we determined that the gene encoding OmpU in toxigenic strains potentially encodes VAPs and exhibits allelic diversity in environmental populations [27]. Interestingly, unlike the *ompU* allele encoded by toxigenic strains, the allele from the environmental isolate *V. cholerae* GBE1114 does not confer virulence-adaptive properties such as bile resistance and leads to impaired intestinal colonization [27]. We hypothesize that specific VAP-containing domains within the toxigenic *ompU* allele confer these emergent virulence-adaptive properties.

Given that these traits are contained within the variable regions of these two divergent alleles, in this article, we reasoned that these natural allelic variations could also provide us with an ideal model system to identify the specific genetic variations that contribute to AMR and the emergence of virulence-adaptive traits. A preliminary sequence survey exposes large differences between the toxigenic and the GBE1114 alleles making it unfeasible to select any particular set of residues (74.7% identity). To narrow down the potential regions associated with the emergence of AMR, first, we examined the natural variability of *ompU* uncovering 41 alleles of the porin out of over 1600 sequences analyzed. These alleles form two phylogenetic clusters with distinct evolutionary paths and striking diversity. We selected 7 representative alleles from each cluster encompassing the diversity of *ompU* and constructed 14 isogenic mutants where we exchanged the toxigenic allele for an environmental one. We exposed the *ompU* mutant strains to host antimicrobials (e.g. bile) to determine how genotypic variations affect phenotypic outcomes. Conservation and structural analyses of each residue within the alleles from strains that exhibit resistant phenotypes versus sensitive ones led to the identification and characterization of four conserved domains among alleles that conferred bile resistance. Interestingly, a mutant strain where we exchanged the four specific domains from the sensitive allele into the clinical one exhibit a phenotype similar to an *ompU* deletion strain in the presence of bile and the host antimicrobial peptide, P2. Finally, to determine novel potential roles of OmpU, we performed a high-throughput screening using phenotypic microarrays and compared the differential survival and growth of *V. cholerae* N16961 (WT) and a Δ*ompU* mutant. We identified clinically relevant antibiotics where resistance is associated with OmpU. Overall, our assessment of natural allelic variations uncovered functional protein domains associated with AMR, shedding light on the evolutionary processes leading to the emergence of this phenomenon.

## RESULTS

### *ompU* forms two major phylogenetic clusters

OmpU plays a crucial role in several virulence-associated traits in *V. cholerae* such as bile tolerance, antimicrobial peptide resistance, and intestinal colonization [30, 31, 33]. We recently determined that only some alleles of *ompU* confer these observed phenotypes [27]. In the presence of 0.4% bile, *V. cholerae* N16961 (WT) exhibits an average 25% survival as measured by c.f.u. post-treatment, while a Δ*ompU* mutant strain shows only 0.16% survival (p<0.0001) [27]. An isogenic *V. cholerae* N16961 mutant strain encoding the *ompU* allele from the environmental strain *V. cholerae* GBE1114 exhibits 1.4% survival (p=0.02 versus Δ*ompU*) [27]. This allele-dependent phenotype indicates the presence of unique functional residues in the clinical allele of OmpU leading to the emergence of bile resistance. To identify these functional residues, we first performed a sequence alignment of the N16961 (resistant) and the GBE1114 (sensitive) proteins (**Fig 1A**). The protein sequences share 74.7% identity with a plethora of mutations including 3 insertions ranging from 1-11aa per event, 8 deletions, and 30 non-synonymous mutations, relative to WT, overall precluding us from directly identifying potential candidates within clinical OmpU leading to AMR.

**Fig. 1.**
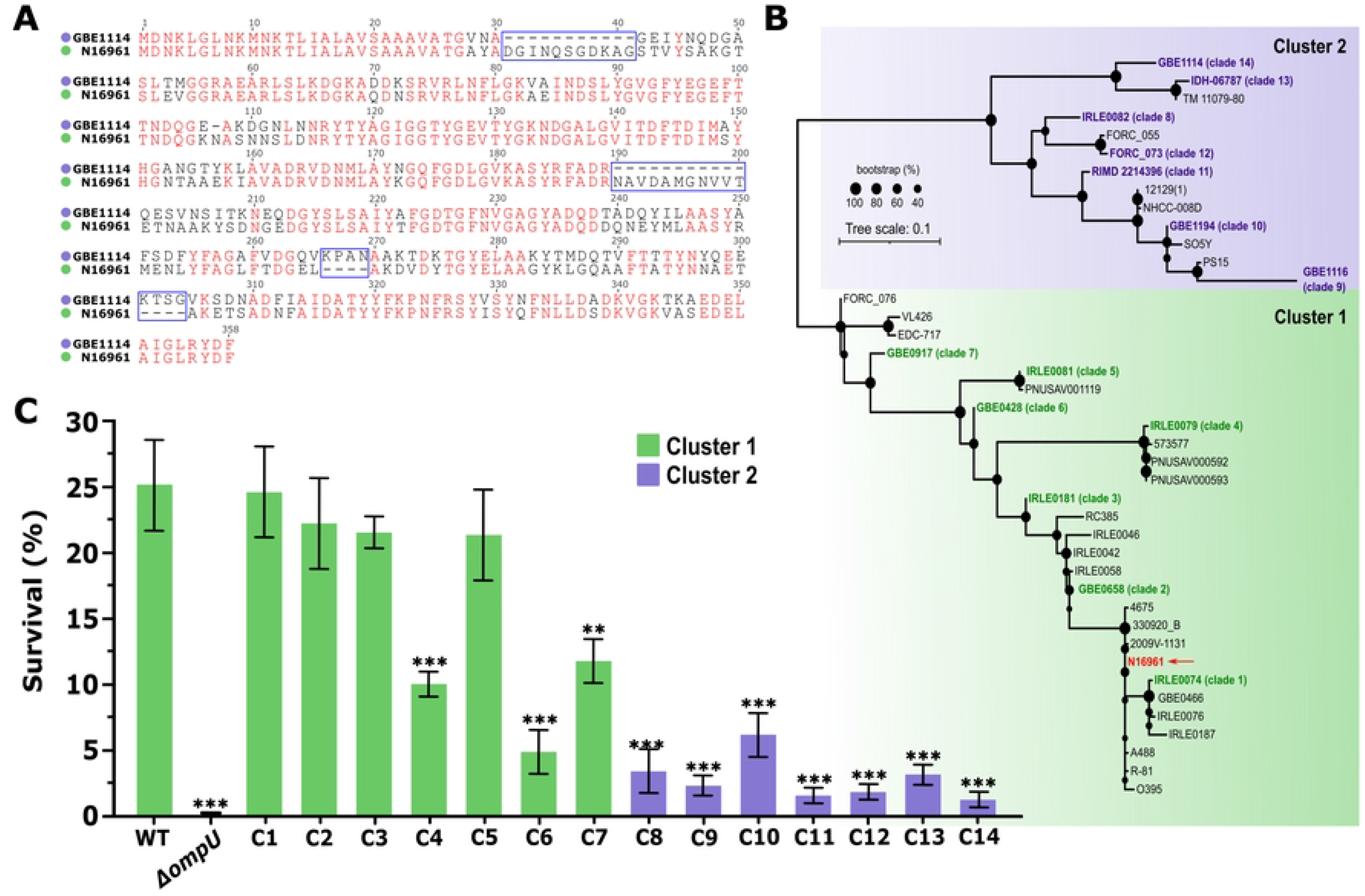
OmpU forms two highly diverse phylogenetic clusters. (A) Sequence alignment of the *ompU* alleles from strains *V. cholerae* N16961 and GBE1114. (B) OmpU has diverged into two major phylogenetic clusters. Red arrow indicates the clinical allele from N16961. Green and purple shading represent Cluster 1 and 2 respectively. (C) Survival of *V. cholerae* mutant strains encoding representative *ompU* alleles in the presence of 0.4% whole bile. Despite the large genetic differences between the alleles, strain survival in bile leads to convergent phenotypes. Error bars represent the standard deviation of the mean from at least three (*N*≥3) independent replicates. Statistical comparisons were performed using Student’s *t*-test and all constructs were compared to the WT unless otherwise stated. \*\**p*<0.01, \*\*\**p*<0.001.

To determine the diversity of OmpU and subsequently correlate it with AMR, we obtained all sequences of the *ompU* gene in *V. cholerae* genomes from public databases, including environmental strains of *V. cholerae* in our collection [27, 29, 34]. *V. cholerae* N16961 OmpU was used as a reference to search for all homologs obtaining a total of 1620 sequences. After a previous clustering of the sequences (100% identity), to eliminate the clonality bias of *V. cholerae*, we generated a phylogenetic tree to examine the evolutionary history of OmpU (**Fig 1B**). The analysis reveals two distinct major clusters composed of 28 clades (Cluster 1; C1) and 13 clades (Cluster 2; C2), respectively (**Fig 1B**). Alleles from toxigenic strains of *V. cholerae* (clinical allele) were confined to one clade within the larger cluster C1. Given this allelic variability, we hypothesized that we could compare the phenotype associated with different alleles and infer genotype-phenotype correlations to discern the functional residues that differentiate the resistant from the sensitive ones. To make this approach tractable we reduced the number of alleles tested to 14 by prioritizing distribution of clades and genotype patterns based on both alignment and phylogeny (**Table 1**). Based on these criteria, we selected 7 alleles within C1, with nucleotide identities ranging from 96% (closest to clinical; IRLE0074) to 75% (furthest; IRLE0079), strengthening the scenario in which the clinical allele of *ompU* emerged as a preadaptation in *V. cholerae* populations prior to host colonization [27, 35]. On the other hand, the nucleotide identities of alleles from C2 range from 72% (closest to clinical; IRLE0082) to 69% (furthest; GBE1116). The environmental allele GBE1114, which is sensitive to bile is found in C2. In comparison to GBE1114, the nucleotide identities of alleles present in C2 range from 89% (closest to GBE1114) to 79% (furthest). From C1 we included the *ompU* alleles from the following *V. cholerae* strains (number between brackets indicates clade number): IRLE0074 (1), GBE0658 (2), IRLE0181 (3), IRLE0079 (4), IRLE0081 (5), GBE0428 (6), and GBE0917 (7), and from C2 IRLE0082 (8) GBE1116 (9), GBE1194 (10), RIMD 2214396 (11), FORC 073 (12), IDH-06787 (13), and GBE1114 (14) (**Table 1, S1 Fig, S1 Appendix**).

**Table 1.**
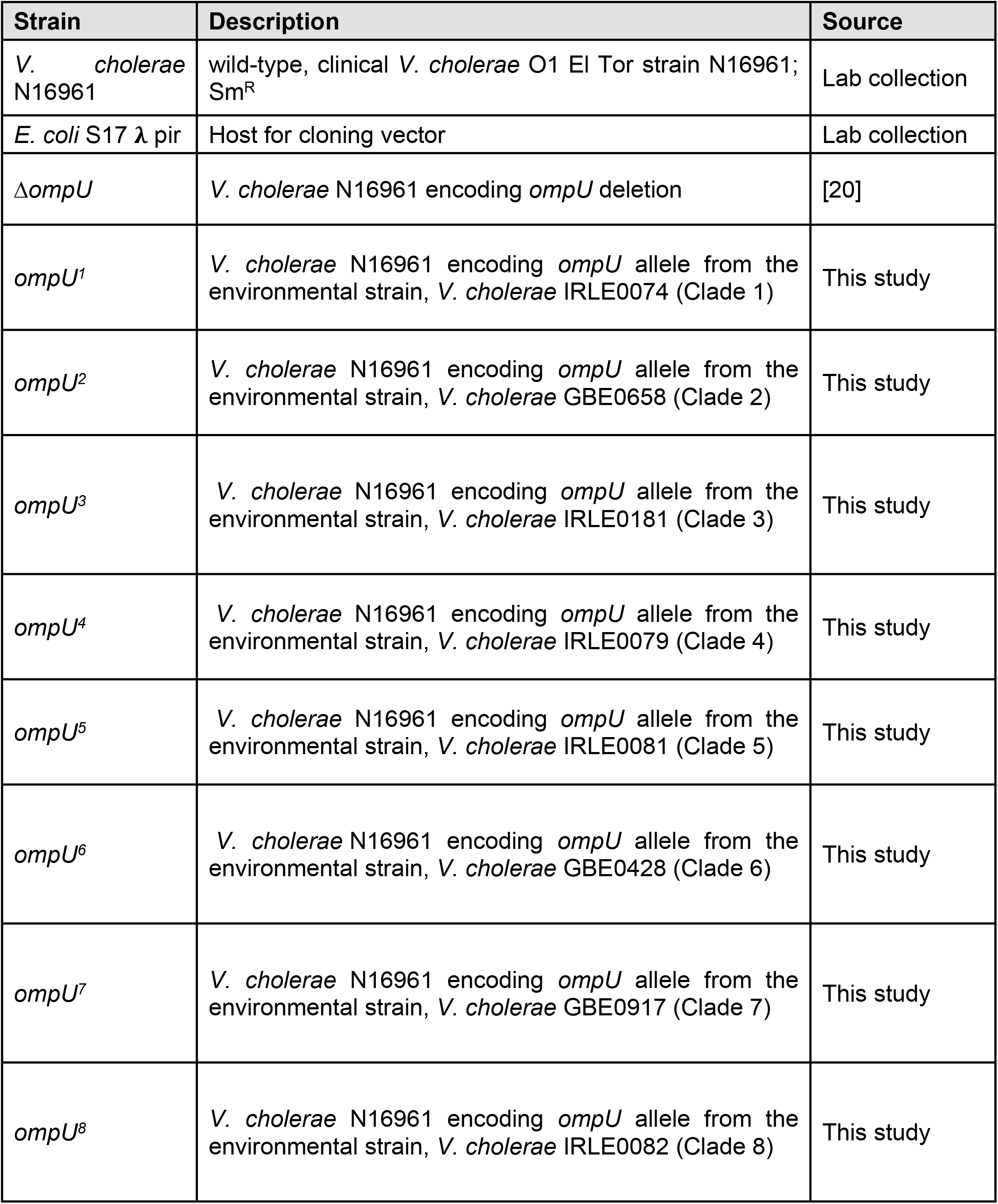
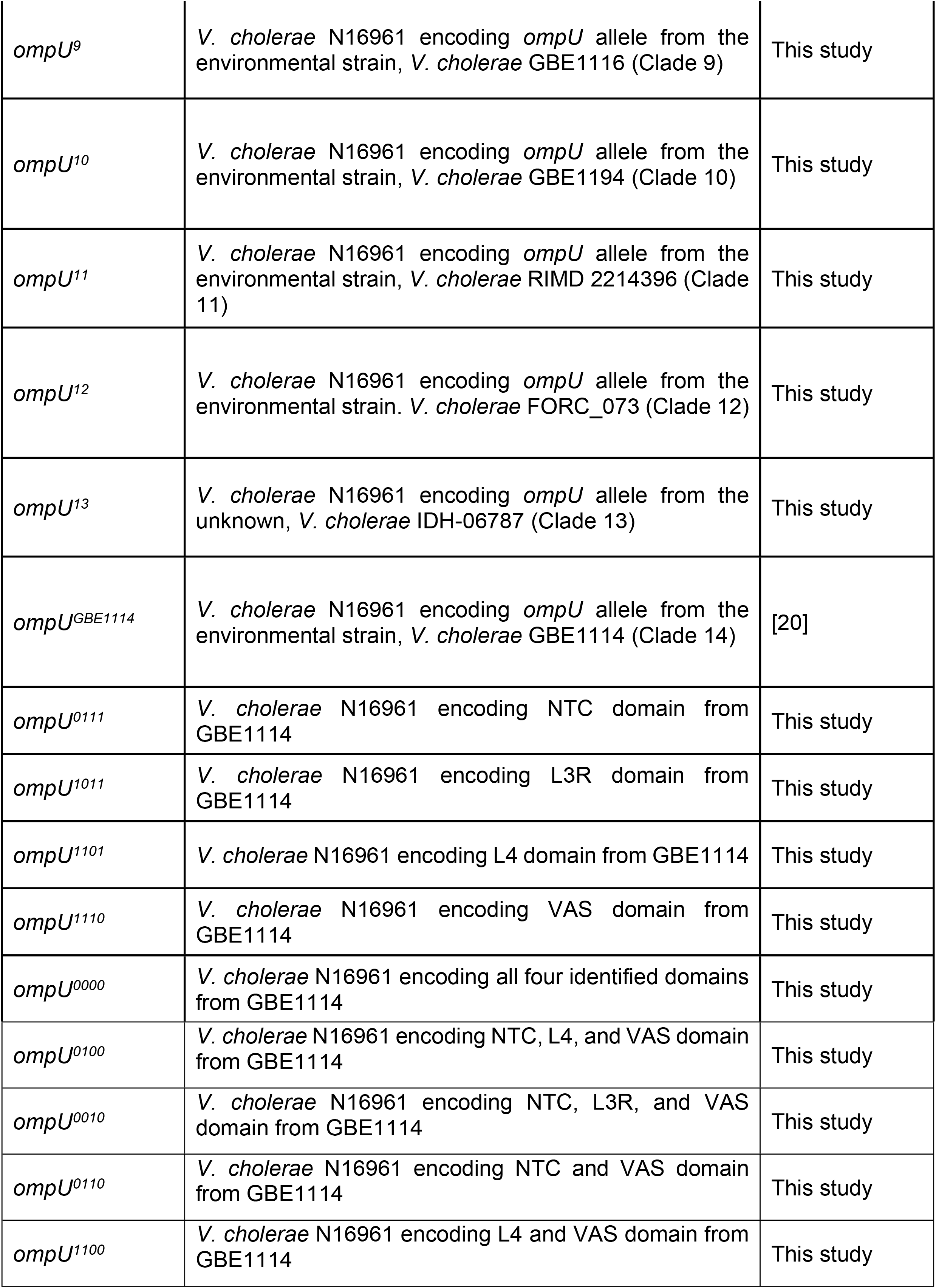
Strains used in this study

| Strain | Description | Source |
| --- | --- | --- |
| *V. cholerae* N16961 | wild-type, clinical *V. cholerae* O1 El Tor strain N16961; Sm<sup>R</sup> | Lab collection |
| *E. coli* S17 $\lambda$ pir | Host for cloning vector | Lab collection |
| $\Delta ompU$ | *V. cholerae* N16961 encoding *ompU* deletion | [20] |
| *ompU*<sup>1</sup> | *V. cholerae* N16961 encoding *ompU* allele from the environmental strain, *V. cholerae* IRLE0074 (Clade 1) | This study |
| *ompU*<sup>2</sup> | *V. cholerae* N16961 encoding *ompU* allele from the environmental strain, *V. cholerae* GBE0658 (Clade 2) | This study |
| *ompU*<sup>3</sup> | *V. cholerae* N16961 encoding *ompU* allele from the environmental strain, *V. cholerae* IRLE0181 (Clade 3) | This study |
| *ompU*<sup>4</sup> | *V. cholerae* N16961 encoding *ompU* allele from the environmental strain, *V. cholerae* IRLE0079 (Clade 4) | This study |
| *ompU*<sup>5</sup> | *V. cholerae* N16961 encoding *ompU* allele from the environmental strain, *V. cholerae* IRLE0081 (Clade 5) | This study |
| *ompU*<sup>6</sup> | *V. cholerae* N16961 encoding *ompU* allele from the environmental strain, *V. cholerae* GBE0428 (Clade 6) | This study |
| *ompU*<sup>7</sup> | *V. cholerae* N16961 encoding *ompU* allele from the environmental strain, *V. cholerae* GBE0917 (Clade 7) | This study |
| *ompU*<sup>8</sup> | *V. cholerae* N16961 encoding *ompU* allele from the environmental strain, *V. cholerae* IRLE0082 (Clade 8) | This study |
| *ompU*<sup>9</sup> | *V. cholerae* N16961 encoding *ompU* allele from the environmental strain, *V. cholerae* GBE1116 (Clade 9) | This study |
| *ompU*<sup>10</sup> | *V. cholerae* N16961 encoding *ompU* allele from the environmental strain, *V. cholerae* GBE1194 (Clade 10) | This study |
| *ompU*<sup>11</sup> | *V. cholerae* N16961 encoding *ompU* allele from the environmental strain, *V. cholerae* RIMD 2214396 (Clade 11) | This study |
| *ompU*<sup>12</sup> | *V. cholerae* N16961 encoding *ompU* allele from the environmental strain. *V. cholerae* FORC_073 (Clade 12) | This study |
| *ompU*<sup>13</sup> | *V. cholerae* N16961 encoding *ompU* allele from the unknown, *V. cholerae* IDH-06787 (Clade 13) | This study |
| *ompU*<sup>GBE1114</sup> | *V. cholerae* N16961 encoding *ompU* allele from the environmental strain, *V. cholerae* GBE1114 (Clade 14) | [20] |
| *ompU*<sup>0111</sup> | *V. cholerae* N16961 encoding NTC domain from GBE1114 | This study |
| *ompU*<sup>1011</sup> | *V. cholerae* N16961 encoding L3R domain from GBE1114 | This study |
| *ompU*<sup>1101</sup> | *V. cholerae* N16961 encoding L4 domain from GBE1114 | This study |
| *ompU*<sup>1110</sup> | *V. cholerae* N16961 encoding VAS domain from GBE1114 | This study |
| *ompU*<sup>0000</sup> | *V. cholerae* N16961 encoding all four identified domains from GBE1114 | This study |
| *ompU*<sup>0100</sup> | *V. cholerae* N16961 encoding NTC, L4, and VAS domain from GBE1114 | This study |
| *ompU*<sup>0010</sup> | *V. cholerae* N16961 encoding NTC, L3R, and VAS domain from GBE1114 | This study |
| *ompU*<sup>0110</sup> | *V. cholerae* N16961 encoding NTC and VAS domain from GBE1114 | This study |
| *ompU*<sup>1100</sup> | *V. cholerae* N16961 encoding L4 and VAS domain from GBE1114 | This study |

### Naturally occurring allelic variations of OmpU lead to phenotypic diversity in the presence of bile

To examine the phenotypes associated with the different representative *ompU* alleles, we constructed 14 isogenic mutants, each encoding one of them, in the genomic background of *V. cholerae* N16961 (**Table 1**). First OmpU protein stability was verified for each strain using immunoblots (**S2 Fig**). Subsequently, we examined the resistance of these strains in the presence of bile (**Fig 1C**). We observed that the overall resistance pattern in the presence of bile broadly correlates with tree topology, with several strains from C1 exhibiting a resistance phenotype like WT whereas the ones from C2 (closer to GBE1114) show a sensitive one (**Fig 1C**). Specifically, strains *ompU*^*1*^, *ompU*^*2*^, *ompU*^*3*^, and *ompU*^*5*^ from C1 display similar resistance to bile as the WT strain. Interestingly, *ompU*^*4*^, *ompU*^*6*^ and *ompU*^*7*^ warrant some further investigations as they show a decrease in survival in the presence of bile, with *ompU*^*6*^ exhibiting a phenotype similar to some strains from C2 (*ompU*^*8*^ and *ompU*^*10*^) (**Fig 1C**). Strains encoding alleles from C2 exhibit on average ∼1-log decrease in survival compared to the WT strain (**Fig 1C**). Those encoding alleles from clades 9, 11 and 12 are the most sensitive to bile when compared to WT and their survival is similar to the strain encoding the allele from GBE1114 (clade 14).

### Genotype to phenotype associations uncover potential domains associated with bile resistance

Phenotypic characterization of *ompU* alleles reveal conserved resistance patterns stemming from diverse genotypes (**Fig 1**). To determine functional domains unique to alleles that confer resistance to bile (resistant alleles), we first created groups of resistant and sensitive alleles based on statistical significance. Using this criterion, we generated two groups with distinct phenotypes: **A)** resistant (*ompU*^*1*^, *ompU*^*2*^, *ompU*^*3*^, and *ompU*^*5*^): alleles that displayed no statistically significant difference in survival when compared to WT and **B)** sensitive (*ompU*^*9*^, *ompU*^*11*^, and *ompU*^*12*^): alleles that displayed no statistically significant difference in survival when compared to GBE1114 (clade 14). These two groups with similar phenotypes but diverse genotypes provided an ideal tool to identify the functional residues contributing to bile resistance in clinical OmpU. We performed a Scorecons analysis of the resistant and sensitive groups to assist the identification of potential functional residues unique to resistant alleles. Based on our analysis, we identified a total of 244/350 conserved positions at the protein level (=1) and 106 positions where residues were not conserved (<1). Of the 106 variable positions, 82 (75%) were distributed in large groups (≥4 or more consecutive variable or inserted residues) within external loops or internal modules of the porin based on published annotations [36, 37]. Conversely, the remaining 24 (25%) were mainly single amino acid changes or variable groups distributed throughout the hydrophobic transmembrane ß-sheets embedded within the outer membrane. The distribution of these large variable groups of residues in external modules of OmpU further suggests that environmental pressures play a role in selecting emergent adaptive mutations to tolerate adverse conditions. To increase the probability of identifying functional domains conferring resistance to host antimicrobials, we focused primarily on amino acid positions where residue variability occurred in clusters. Based on this parameter, we identified seven amino acid (aa) regions of interest **(S1 Fig)**: 1) 31-43, 2) 102-114, 3) 153-158, 4) 193-203, 5) 264-272, 6) 293-305 and 7) 336-338 relative to the WT allele.

After examining each aligned column and narrowing down to clusters of non-conserved amino acids, we then sought to identify clusters containing changes unique to resistant alleles. Using the criteria described in the materials and methods section, we examined the seven identified regions and of these, regions 1 (31-43aa), and 3 (153-158aa) showed a clear discrimination between the sensitive and resistant alleles. Conservation scores for regions 4 (193-203) and 7 (336-338) were variable for both resistant and sensitive alleles, however, based on the criteria outlined, we observed a trend towards non-conservative for this region with a higher number of nonsynonymous residues occurring in sensitive alleles. Regions 2 (102-114aa), 5 (264-272aa) and 6 (293-305aa) have positions that are so variable that no pattern could be identified even for resistant alleles limiting their relevance as potential domains associated with the emergence antimicrobial resistance in OmpU. Thus, they were excluded from further analysis. Overall, our systematic genotype to phenotype association and comparative sequence analyses led us to identify regions 1, 3, 4 and 7, which we will term N-terminal coil (NTC), L3 region (L3R), L4-loop (L4) and VAS respectively, for subsequent analyses (**Fig 2A, S3 Fig**).

**Fig. 2.**
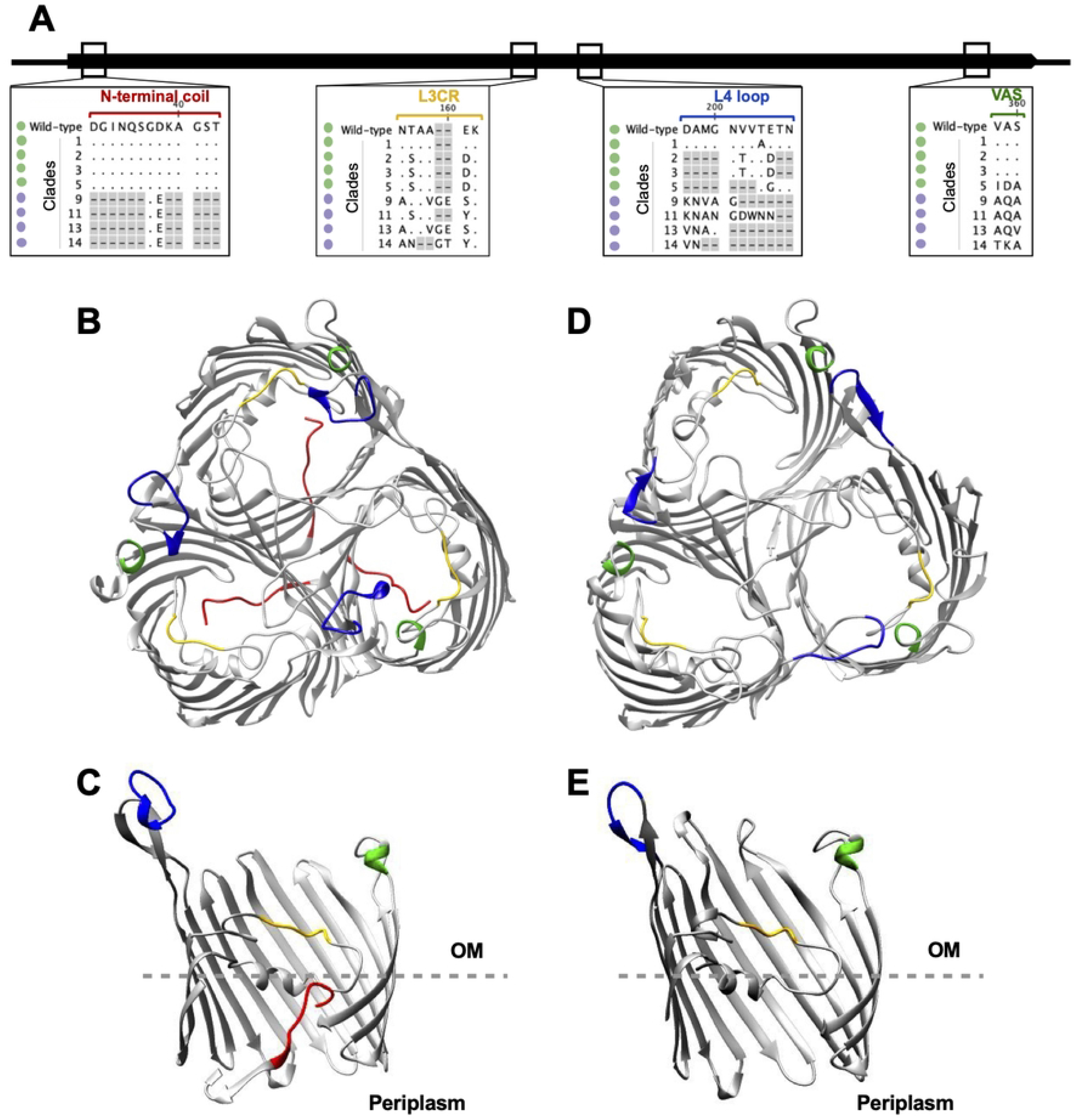
Comparative visualization of domains within OmpU associated with bile resistant phenotypes. (A). Multiple sequence alignment of *ompU* alleles resistant and sensitive to whole bile. Conserved residues are indicated by dots and those absent are indicated by gray boxes. (B-E) Comparative architecture of OmpU domains associated with the bile resistant phenotype. Domains are color coded and visualized in OmpU N16961 and GBE1114. Top and slabbed view of (B, C) N16961 and (D, E) GBE1114, respectively. The identified domains are colored as follows: NTC (red), L3R (yellow), L4 (blue) and VAS (green).

### 3D visualization and structural analysis of identified protein domains

OmpU is a homotrimer that is distributed throughout 60% of the outer membrane [38]. Each monomer consists of 16 ß-sheets embedded within the outer membrane connected by 8 short turns on the periplasmic side and 8 loops on the external side [36] (**Fig 2B**). From the extracellular side, there are eight loops, termed L1-8, connecting these ß-sheets. Of these, six extend into the extracellular space (L1, L4-8) and are exposed to fluctuating environmental conditions, one (L2) overlaps its neighboring monomer and facilitates stabilization of the trimer and the last (L3) folds into the porin and forms a constriction region across the porin that interacts with amino acids on the inner barrel wall [36, 37] (**Fig 2B**). On the periplasmic side of the porin, at the N-terminal region, a coil extends inward into the porin and interacts with L3 [37]. We examined whether the four identified domains translate into structural differences in OmpU using the protein visualization software Chimera [39]. In order to do this, we generated a predicted OmpU protein model of the GBE1114 allele and examined individual and superimposed OmpU protein models of N16961 (OmpU^N16961^) and GBE1114 (OmpU^GBE1114^). As identified in our analyses above, structural comparisons reveal that the NTC domain that pushes inward into OmpU^N16961^ [40] is absent from OmpU^GBE1114^ (**Fig 2B and 2D**). Secondly, amino acid residues present in the L3R domain identified above, are located on the inward L3 loop and differ in charge and hydrophobicity between the two alleles (**Figs 2C and 2E**). OmpU^N16961^ encodes charged residues (E, and K), whereas OmpU^GBE1114^ does not encode any charged residues (**Fig 2A**). Lastly, domains L4R and VAS are located on loops L4 and L8 respectively that protrude into the extracellular space (**Figs 2C and 2E**). Interestingly, overlay of both porins reveal an extended L4 in OmpU^N16961^ when compared to OmpU^GBE1114^. Additionally, the VAS domain is hydrophobic in OmpU^N16961^ whereas in OmpU^GBE1114^ it is basic (**Fig 2A**). Overall, the identified domains translate to several structural differences that may alter protein-protein interactions, modulate open and closed conformation states, or modify overall stabilization of the porin in unfavorable conditions.

### OmpU-mediated resistance to bile requires specific domains present in the clinical allele

To determine whether the domains we identified above play a role in OmpU-mediated AMR, we generated four individual mutant constructs that encode the GBE1114 sequence of each identified domain in the background of *V. cholerae* N16961. For ease of classification, each mutant was assigned a binary code based on the domains encoded, where ‘1’ represents the WT domain from N16961 and ‘0’ represents the domain from GBE1114. This was done for each domain and the strain classification is as follows: *ompU*^*0111*^ (NTC), *ompU*^*1011*^ (L3R), *ompU*^*1101*^ (L4) and *ompU*^*1110*^ (VAS). In addition, to determine whether a synergistic effect exists among the domains, we constructed an isogenic mutant strain encoding an *ompU* allele with each of the four domains exchanged with the corresponding sequence from GBE1114. This allele was assigned the binary code ‘0000’. First, we verified protein stability using immunoblots and subsequently examined the survival of the mutants in the presence of 0.4% whole bile (**Fig 3A and B**). Interestingly, two individual domain mutants, *ompU*^*0111*^ (NTC) and *ompU*^*1011*^ (L3R), show the greatest susceptibility to bile with survival rates of 9.6% and 4.5% respectively (**Fig 3B**). The *ompU*^*1101*^ mutant exhibits 17% survival rate, whereas the *ompU*^*1110*^ mutant shows no survival defect, and displays similar survival as the WT. The *ompU*^*0000*^ “null” mutant has an average survival rate of 3.4%, closely resembling the phenotype of the strain encoding GBE1114 (**Fig 3B**). Interestingly, the *ompU*^*0000*^ mutant shows similar survival in the presence of bile to the single domain mutant *ompU*^*1011*^. Overall, our results indicate that, independently, the NTC and L3R domains contribute to the OmpU ability to confer bile resistance.

**Fig. 3.**
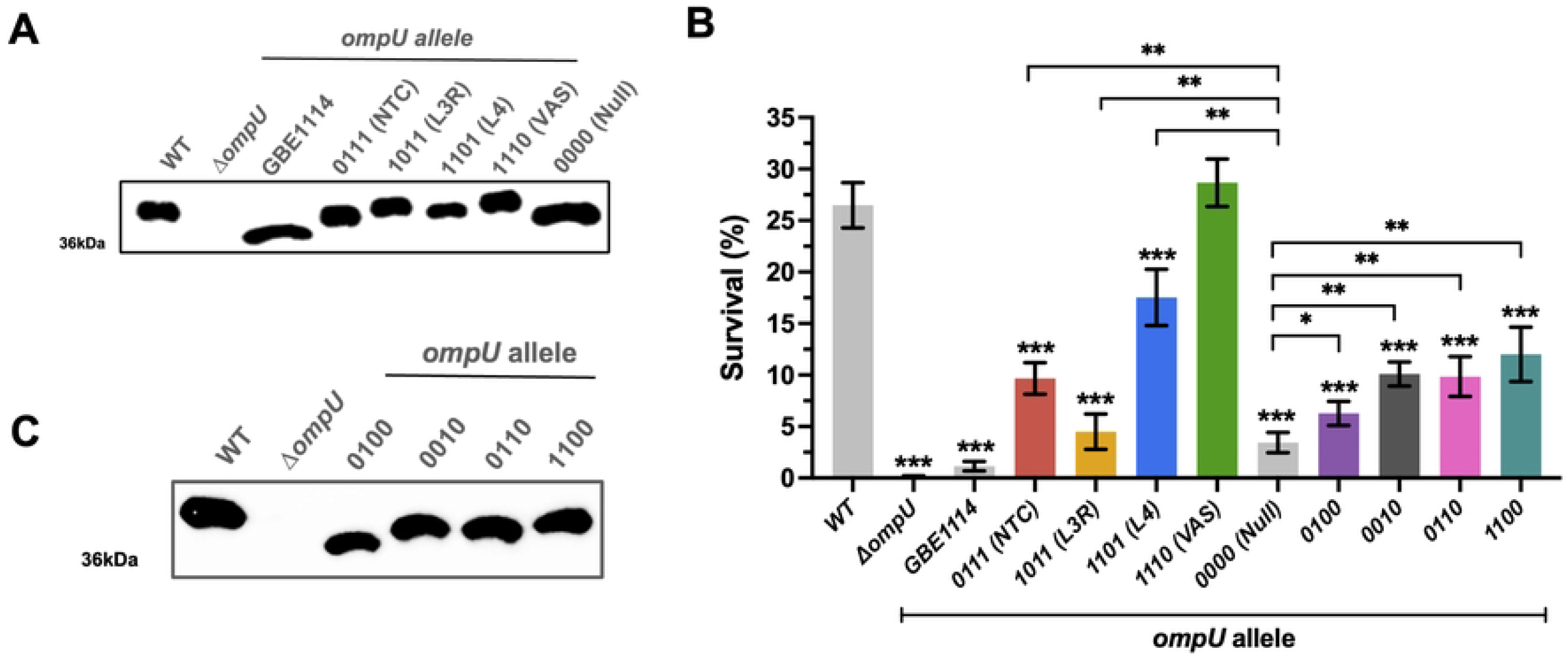
OmpU domain mutants exhibit differential resistance to 0.4% bile. (A) Immunoblot analysis of individual OmpU domain mutants. (B) Survival of individual and naturally occurring *ompU* domain combination mutants in the presence of 0.4% whole bile. (C) Immunoblot analysis of naturally occurring OmpU domain mutants. Error bars represent the standard deviation of the mean from at least three (*N*≥3) independent replicates. Statistical comparisons were performed using Student’s *t*-test and all constructs were compared to the WT unless otherwise stated. \**p*<0.05, \*\**p*<0.01, \*\*\**p*<0.001.

Next, we examined the diversity of naturally-occurring domain combinations in our representative dataset of *ompU* alleles to unearth potential synergistic effects between the domains. We identified four naturally occurring domain combinations and classified each using the established binary nomenclature. The domains combinations are as follows: *ompU*^*0100*^ (GBE0917), *ompU*^*0010*^ (IDH-06787), *ompU*^*0110*^ (GBE1116), and *ompU*^*1100*^ (IRLE0081). Next, as described above, we constructed mutant strains encoding those domain combinations and tested their bile resistance profiles after testing the stability of OmpU in those mutants (**Fig 3C**). Upon exposure to 0.4% whole bile, all of the mutants exhibit a decrease in bile resistance with survival ranging from 6 to 12% (**Fig 3B**). While we determined that L3R is crucial for resistance to bile, the mutant *ompU*^*0100*^ reveals that presence of only the clinical L3R domain cannot recover the bile-resistant phenotype, further suggesting a synergistic effect between the domains. Secondly, strains encoding alleles with the clinical L4 domain (*ompU*^*0010*^) or the L3R and L4 domains (*ompU*^*0110*^), display a minor increase in survival when compared with *ompU*^*0100*^ (**Fig 3C**). Lastly, the strain encoding both the clinical NTC and L3R domain (*ompU*^*1100*^) shows the highest resistance to bile (12%) when compared to other domain combinations (**Fig 3C**). Our results indicate that individually both NTC and L3R domains are crucial for increased resistance to bile (**Fig 3C)**. However, the synergistic effect of all domains appears to be essential for AMR in the clinical OmpU to emerge.

### OmpU domain mutants exhibit differential survival upon exposure to various antimicrobials

OmpU in toxigenic *V. cholerae* is known to confer resistance to the host antimicrobial peptide P2, polymyxin B (PB), and tolerance to organic acid [32, 33, 41]. To determine whether these OmpU-associated phenotypes are allelic- and domain-dependent, we phenotypically characterized the *ompU*^*GBE1114*^ and *ompU*^*0000*^ domain mutants in the presence of the cationic antimicrobial peptides (CAMPs) P2 and PB, and organic acid.

#### Antimicrobial peptide P2

After exposure to P2, WT has a 91% survival rate whereas the Δ*ompU* mutant exhibits a survival rate of 7.5%, comparable to previously published studies [33]. Interestingly, the mutant *ompU*^*GBE1114*^ shows 12% survival, and the domain mutant *ompU*^*0000*^ 23%. (**Fig 4A**). All these strains exhibit a statistically significant difference in survival when compared to WT (*p<*0.001), indicating that survival in the presence of P2 is both allelic and domain-dependent (**Fig 4A**). Next, we determined the role of the individual domains in resistance to P2. Interestingly, we found that all the domain mutants show a decrease in survival ranging between 43-52% compared to WT (p<0.0001) (**Fig 4**). A potential synergistic effect between the domains might explain their difference with *ompU*^*0000*^.

**Fig. 4.**
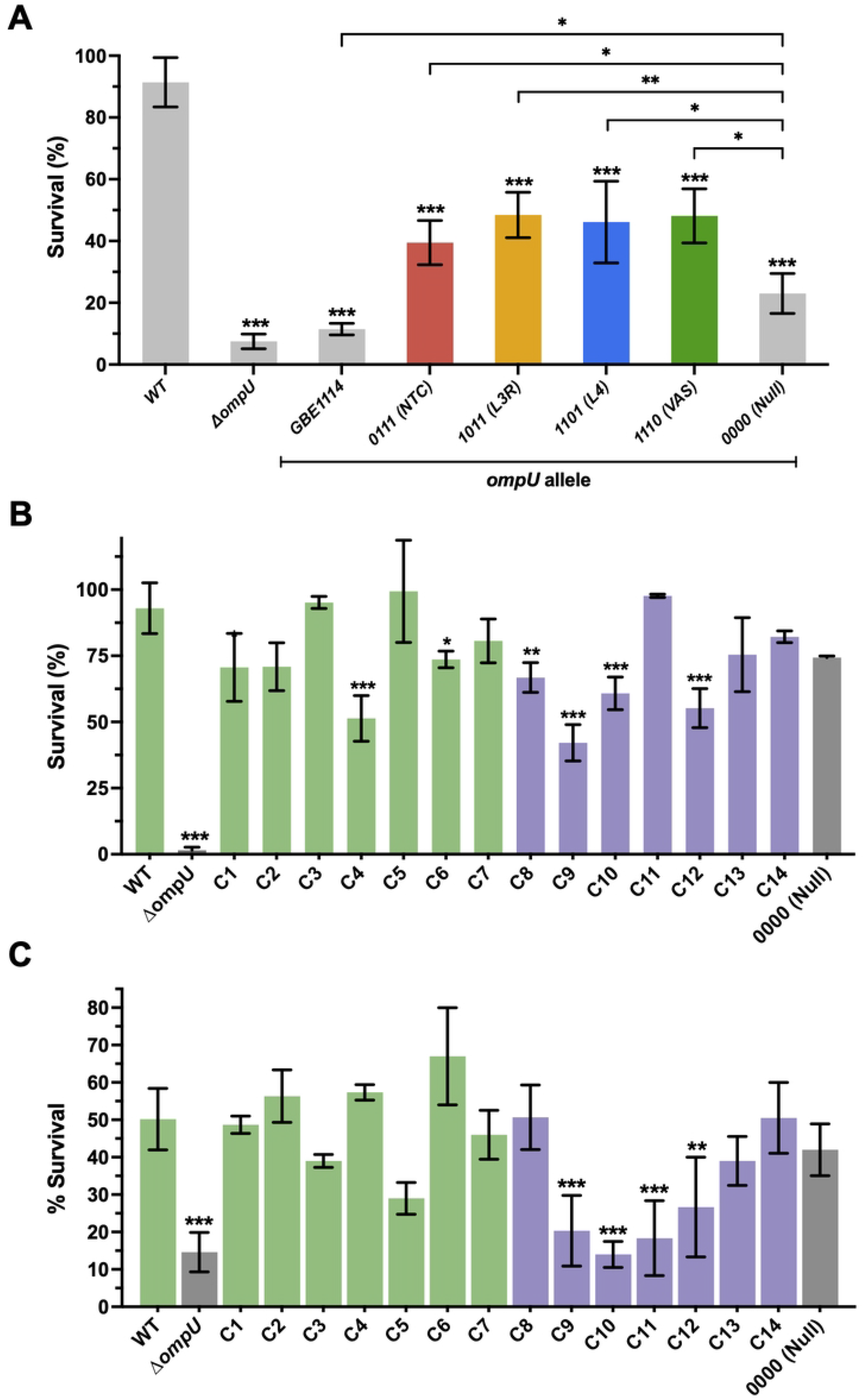
Role of protein domains in OmpU-associated phenotypes. Survival of *V. cholerae* N16961 *ompU* mutant strains in the presence of (A) 200ug P2, (B) polymyxin B (450UmL^−1^) and (C) organic acid (0.005X). Error bars represent the standard deviation of the mean from at least three (*N*≥3) independent replicates. Statistical comparisons were performed using Student’s *t*-test and all constructs were compared to the WT unless otherwise stated. \**p*<0.05, \*\**p*<0.01, \*\*\**p*<0.001, \*\*\*\**p*<0.0001.

#### Polymyxin B

Secondly, in the presence of PB, as previously determined, Δ*ompU* show a drastic decrease in survival (11.3%) compared to WT (92.9%). Unlike for P2, neither *ompU*^*GBE1114*^ nor *ompU*^*0000*^ show a change in survival compared to WT (**Fig 4B**). As shown in Fig 1C, there is phenotypic diversity in the alleles tested in the presence of bile, thus, we considered that it would be possible that resistance to PB might be allelic-dependent yet not associated with the specific allele from strain GBE1114. In order to examine this, we exposed the other 13 mutant strains encoding the different *ompU* alleles to PB and measured their survival (**Fig 4B**). Overall, we did not find major differences between the mutant strains and the WT, certainly none of them resembling the Δ*ompU* mutant (**Fig 4B**). Strains with the *ompU* allele from clades 4, 9, and 12 show the lowest survival in the presence of PB (between 42% and 55%) but not enough to warrant strong evidence that PB resistance is allelic-dependent. Furthermore, there is no clear pattern that associates these strains with a given cluster.

#### Organic acid

On the other hand, even though neither *ompU*^*GBE1114*^ (C14) nor *ompU*^*0000*^ show a decrease in survival compared to WT when cells were exposed to acidic pH, the role of OmpU in acid tolerance appears to be allelic-dependent (**Fig 4C**). Specifically, strains from clades 9, 10 and 11 show between 14% and 20% survival in acidic pH, similar to Δ*ompU* (12%), whereas the WT show an average survival of 50% (**Fig 4C**). All the strains with increased sensitivity to organic acid belong to Cluster 2. Overall, our results uncover complex associations between the different alleles and domains of *ompU* and their ability to confer phenotypes associated with resistance to host antimicrobials.

### OmpU confers resistance to Rifamycin SV

OmpU is ubiquitous in the outer membrane of *V. cholerae* and, as our results indicate, appears to have more nuances than expected in the form of allele-associated domains [27]. Based on this, we considered that it is possible that the porin plays other roles in *V. cholerae* survival beyond those already known. In order to test this, we screened for potential novel functions of OmpU by performing high-throughput assays using phenotypic microarrays (**Fig 5A**). We tested for sensitivity to chemicals and antimicrobials (960 compounds and conditions), growth and survival at different pH (96 conditions) and osmolytes (96 conditions). We compared the growth of the WT and Δ*ompU* mutant strains under these conditions and identified thirteen antimicrobial compounds in the presence of which the WT strain exhibits at least two-fold greater area under the curve (AUC) compared to the Δ*ompU* mutant (**Fig 5A**). Of particular interest, four of these compounds are antibiotics that have been used in clinical settings: rifamycin, oxytetracycline, cinoxacin and troleandomycin [42-46]. Subsequently, we performed *in-vitro* survival assays in the presence of these antibiotics to corroborate the OmpU-dependent phenotypes observed in the phenotypic microarrays and determine whether resistance to these compounds is allelic-dependent. Troleandomycin was not commercially available and could not be included in our *in vitro* assays. Given that the concentration of antimicrobials provided in the phenotypic microarray plates are proprietary information, first we performed titrations of each antimicrobial to determine if we could replicate the growth difference in a survival assay. Furthermore, to examine the role of allelic variations on survival in the presence of these antimicrobials, a large enough cut-off for the difference between the WT and Δ*ompU* was deemed necessary. For this, we followed a cutoff in which a) the WT displays at least 25% survival under that antibiotic concentration, and b) at that concentration the Δ*ompU* mutant exhibits at least a 5-fold decrease in survival.

**Fig. 5.**
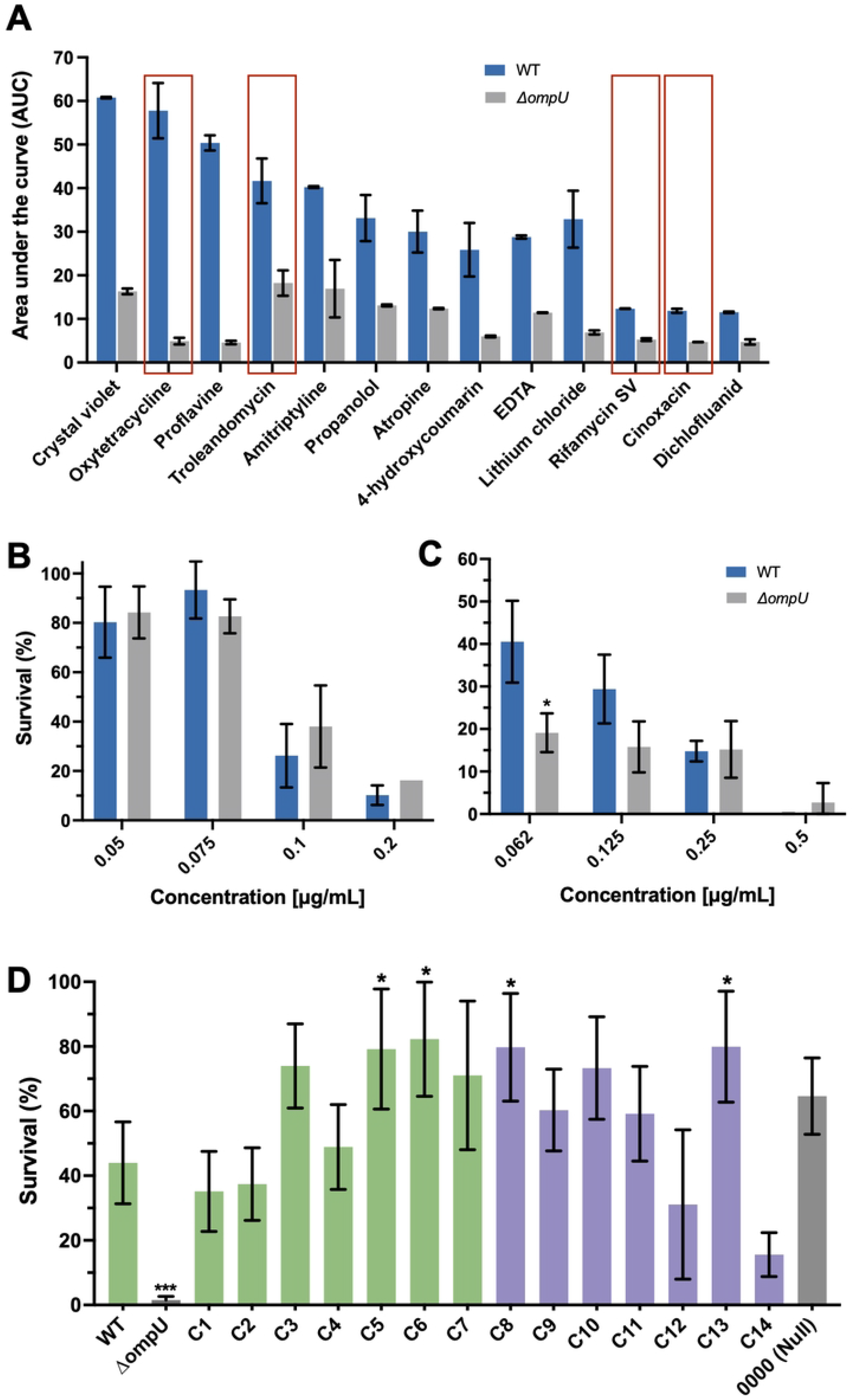
Exploration of novel functions of OmpU. (A) Phenotypic microarrays of *V. cholerae* WT and *ΔompU* strains. Area under the curve (AUC) was calculated to determine the total growth of the strains under exposure to diverse antimicrobial compounds. Only compounds with greater than 2-fold decrease in AUC for Δ*ompU* strain are shown. Red rectangles indicate antibiotics with clinical relevance. All arrays were performed in duplicate. (B, C) Survival of *V. cholerae* N16961 and Δ*ompU* in the presence of varying concentrations of (B) oxytetracycline and (C) cinoxacin. (D) Survival of *ompU* alleles in 7.5μg/mL rifamycin SV. Error bars represent the standard deviation of the mean from at least three (*N*≥3) independent replicates. Statistical comparisons were performed using Student’s *t*-test and all constructs were compared to the WT unless otherwise stated. \**p*<0.05. *N*≥3, *p<0.05.

#### Cinoxacin and oxytetracycline

Based on our titrations of cinoxacin and oxytetracycline, we could not identify a concentration that met our cut-off to infer an OmpU-dependent role in the survival against these antimicrobials (**Fig 5B and C**).

#### Rifamycin SV

First, we tested survival in the presence of rifamycin SV (RSV). The WT strain shows an average survival of 44% and had a survival difference of ∼28-fold when compared to Δ*ompU* (1.5%) (**Fig 5D**). This survival difference provided us with enough potential resolution to observe differences betweem the alleles. As for the conditions tested above (**Fig 4**), next, we examined the survival of the *ompU*^*GBE1114*^ (C14) and *ompU*^*0000*^ strains in the presence of RSV. We found that even though resistance to RSV appears to be allelic-dependent (*ompU*^*GBE1114*^ exhibits decreased survival), the domains that we found associated with bile and P2 survival do not play a role in this phenotype (*ompU*^*0000*^ exhibits WT-level of survival). Our results indicate that other domains within OmpU might be responsible for this phenotype (**Fig 5D**). Finally, we tested the survival of the other *ompU* allele mutant strains in the presence of RSV and found that four of them (*ompU*^*5*^, *ompU*^*6*^, *ompU*^*8*^, and *ompU*^*13*^) exhibit slightly increased resistance to the antibiotic than the WT (**Fig 5D**). Overall, resistance to RSV is OmpU-dependent and no allele other than *ompU*^*GBE1114*^ display lower survival rate than the WT allele.

## DISCUSSION

The development of antimicrobial resistance among bacterial pathogens is unarguably one of the most critical public health threats that we face and fuels the arms race to combat hospital and community-acquired infections [47-49]. Furthermore, this phenomenon is closely linked to the process of pathogen emergence [50-52]. Gram-negative pathogens pose a major obstacle towards tackling this as the presence of their highly selective outer membrane (OM) critically facilitate adaptations for AMR. Porins represent one of the major proteins present in the OM and are often directly implicated in AMR mechanisms. OmpU, a homotrimer porin that is distributed throughout ∼60% of the OM of *V. cholerae*, is directly associated with the emergence of the pathogen and confers resistance to numerous host antimicrobials [38, 53, 54]. Here, using the allelic diversity found in natural populations of *V. cholerae*, we developed a framework that allowed us to identify domains within the porin associated with the emergence and evolution of AMR in pathogenic strains of the bacterium. Examination of the distribution and variability of *ompU* alleles within *V. cholerae* reveals a highly diverse landscape in which two large clusters encompass all the alleles of the porin. This vast diversity of allelic variants suggests a scenario in which variations in this protein are strongly driven by environmental pressures and likely confer a set of phenotypes much wider than anticipated. It is known that OmpU acts as the receptor of phage ICP2, which directly interacts with domains that are present on extracellular loops of the porin (L4 and L8) [55, 56]. Nonetheless, the extensive variability of OmpU goes beyond the region of the porin that acts as the phage receptor. This indicates that, despite phages likely being major drivers in the selection of these variants, the porin must undergo other strong selective pressures that led to this uncanny diversity.

Bile is an anionic salt under the pH conditions present in the small intestine and requires facilitated transport to disrupt the cell integrity of enteric pathogens such as *V. cholerae* [54, 57, 58]. Upon exposure to bile, *V. cholerae* activates a signaling cascade leading to the upregulation of *ompU* and promoting its survival in the SI [30, 31, 57, 59-61]. Our results reveal that the ability of the OmpU allele encoded by toxigenic strains to confer resistance to bile is due to four domains encoded within different regions of the porin. Phenotypic characterization of the four identified domains, reveals that two are the most crucial for bile resistance: the NTC and L3 domains. Protein modelling of the WT allele show that the NTC domain extends from the periplasm inwards into the porin and interacts with the inner loop 3 (L3), which traverses the upper region of the OmpU towards to the extracellular side. Domains analogous to NTC and L3 in other porins contribute to pore size and altered open and close conformation states [37, 62-67]. Given our findings, it is likely that the presence of the N-terminal coil contributes to a smaller pore size [37] in the WT allele leading to exclusion of bile molecules in contrast to the sensitive alleles. In addition, charged residues on the L3 domain in the WT allele may facilitate longer closed confirmation states and prevent bile influx.

Cationic antimicrobial peptides (CAMPs) are natural peptide products that kill bacteria through membrane destabilization [68-70]. CAMPs selectively target Gram-negative pathogens due to their affinity for negatively charged outer membranes [68]. To evade CAMPs such as P2 or polymyxin B, *V. cholerae* activates the envelope stress response in a sigma E-dependent manner [32, 33]. The initiation of this cascade is dependent on the presence of OmpU [32, 33]. Interestingly, our data reveals that OmpU-associated CAMP resistance is allelic and domain dependent only in the presence of the host-derived peptide P2. On the other hand, polymyxin B is an antimicrobial peptide produced by *Paenibacillus polymyxa* [71], a bacterium that can be found in aquatic habitats [72]. It is likely that recurrent exposure to this CAMP led to population bottlenecks that selected for alleles that confer widespread resistance. Given the differences in the observed survival pattern, this difference in mechanism of action as well as the potential role of other genes in counteracting these antimicrobials remains enigmatic and warrants further investigations.

We determined that all individual domains played a role in resistance to P2. However, absence of no individual domain leads to a survival pattern similar to *ompU*^*0000*^ suggesting a synergistic effect among the domains leading to increased sensitivity in *ompU*^*0000*^. Computational modeling suggests that residues on N-terminal coil interact with the YDF domain to facilitate porin stabilization [37]. We hypothesize that this may function to expose the YDF domain upon envelope stress increasing the survival of clinical strains in the presence of host antimicrobials. Furthermore, the extended negatively charged L4 loop may act as an interacting site for positively charged P2 molecules and reduce its translocation into the cell. Also, interactions between charged residues on L3CR and the barrel wall are likely to affect open/close conformation states of the porin. Lastly, regions upstream of the C-terminal YDF domain are crucial for the stabilization of the anti-anti sigma factor RseB. This facilitates the first cleavage step in the envelope stress response signaling cascade, regulated by RpoE [73, 74]. Given these observed differences and the significance of OmpU to AMP resistance, we propose that these structural differences cooperatively contribute to activating RpoE upon exposure to P2. For some phenotypes tested, such as polymyxin B, neither allelic nor domain-dependent pattern was observed. This indicates that in the presence of these compounds, there may be compensatory events that offset its effects. On the other hand, survival in the presence of organic acid is allelic-dependent but appears to require the presence of *ompU* domains that are distinct from the ones required for bile or P2 survival. Future work will focus on elucidating these specific domains and examining how they might influence other functions of OmpU.

Our findings highlight the astonishing nuances of OmpU and its multifaceted role in the emergence and pathogenesis of *V. cholerae*. Adaptations leading to antimicrobial resistance are essential to the continued success of *V. cholerae* as a pathogen. Recent reports stress the critical importance of monitoring AMR in *V. cholerae* as the bacterium is gaining it at unprecedented rates [75-77]. Examination of the natural variability of OmpU has provided us with a unique tool to determine how genotypic variations within a porin can foster emergent phenotypes in bacterial pathogens. To date, the potential downstream regulatory effects associated with the emergence of the identified domains remain unknown. Future work will address this and other critical questions that will take us closer to dissecting the complex AMR phenomenon.

## MATERIALS AND METHODS

### Bacterial strains and culture conditions

Strains used in this study (Table 1) were routinely grown aerobically on Luria Bertani (LB) media at 37 ºC for approximately 16hrs, unless otherwise stated. Antibiotic concentrations used for plasmid or strain selection are as follows, kanamycin (Kn) 45μg/mL, polymyxin B (PB) 50μg/mL, and streptomycin (Sm) 1mg/ml.

### OmpU phylogenetic tree

All available *V. cholerae* genomes were downloaded from the National Center for Biotechnology Information (NCBI, release 244 July 2021). Then, the DNA sequence coding for the outer membrane protein OmpU (VC_0633) in *V. cholerae* O1 biovar El Tor str. N16961 was used as a reference to obtain all the homologous sequences in the genomes using DIAMOND [78]. Sequences were clustered with CD-HIT [79] and aligned using MUSCLE [80]. Maximum-likelihood tree was then constructed using RAxML [81] with the following parameters: “-f a” algorithm, 100 bootstrap replicates, PROTGAMMAJTT model.

### Construction of mutant strains

*V. cholerae* constructs were all created in the N16961 background following previously published methods [27]. Briefly, for allelic exchange of each representative clades of *ompU*, the allele of interest was either PCR amplified from environmental strains in our collection (Table 1) or synthesized (Gene Universal) if strain was unavailable in our collection. In addition, we PCR amplified 500bp upstream of the 5’ end of *V. cholerae* N16961 *ompU* and 500bp downstream of the 3’ end of the gene. PCR or synthesized products were digested and ligated into the suicide vector pKAS154 using a four-segmented ligation. Ligation products were electroporated into *Escherichia coli* S17λpir and the products plated on LB supplemented with Kn to select for plasmid uptake. PCR amplicons of plasmid constructs were sequenced and compared to environmental allele for verification. Strains carrying the insert were conjugated with *V. cholerae* N16961 and selected for using a series of antibiotics outlined here [82]. PCR amplification was performed to screen for the *ompU* allele and potentially positive samples were sequenced and verified by sequence comparison.

### Western Blot analysis

Overnight cultures of the WT and mutant constructs were diluted to OD_600nm_ 1.0, centrifuged, resuspended in 100μL of sample loading buffer and boiled for 5mins. Samples were diluted 1:5 in sample loading buffer and quantified using the Pierce BCA Protein Assay Kit (Thermo Scientific). Lysates were normalized to 100μg/mL in 1X Bolt™ LDS Sample Buffer (Invitrogen). A total of 2 μL of normalized lysates were loaded into precast 12% Tris-glycine gels (Invitrogen) alongside protein standards (SeeBlue Pre-stained Protein Standard, Invitrogen). Gels were run at room temperature for 150 min at 80 V in running buffer. Blots were transferred to nitrocellulose membranes using the iBlot system (Invitrogen). After the transfer, membranes were removed and placed in blocking buffer (TBST supplemented with 5% non-fat milk) and incubated overnight at 4°C. Subsequently, blocking buffer was removed and replaced with TBST buffer supplemented with 1:30,000 anti-OmpU rabbit primary antibody (GenScript) and incubated for 1h. Membranes were washed three times with TBST after which TBST supplemented with 1:10,000 of anti-rabbit secondary antibody (GenScript) was added and incubated for 30min at room temperature. After incubation with the secondary antibody, membranes were washed three times with TBS buffer. Reagents from the Pierce ECL kit were mixed according to manufacturer’s instructions and chemiluminescence was detected using BioRad ChemiDoc XRS system.

### Sequence comparison and evaluation of resistant and sensitive *ompU* alleles

Multiple sequence alignments (S1 Fig) were generated using CLC Genomics workbench 21 and the conservation score for the residues in each aligned column calculated using the entropy-based method, Scorecons [83]. Based on the 7 entropic states in which amino acids exists, we examined the diversity of each residue within an aligned column and designated a score between 0 (not conserved) and 1 (conserved). To determine whether the residues of sensitive alleles influence conservation at each site, we subtracted conservation scores obtained for resistant alleles from those obtained for resistant and sensitive alleles. A value <0 indicates presence of non-synonymous residues at a given site. On the other hand, a site with a score equal to zero (0) would indicate that the residues are either conserved at this site or there is significant number of variations between both groups and would therefore not be of interest given that there is no exclusivity to either group. A score that is >0 would indicate that there is high conservation in the combined conservation comparison but differences in the resistant group. These sites would not be of interest given that the variability does not affect the survival pattern. **Identification of naturally occurring domains**. Based on the four identified domains, each representative allele was examined for changes in conservation score at each domain position when compared to the WT. Based on this criteria, four naturally occurring domain combinations were identified: *ompU*^*0100*^ (GBE0917), *ompU*^*0010*^ (IDH-06787), *ompU*^*0110*^ (GBE1116), and *ompU*^*1100*^ (IRLE0081).

### OmpU structure visualization

Nucleotide sequences of the *ompU* alleles from N16961 and GBE1114 were translated to amino acid sequences using the online tool Expasy and uploaded to SWISS-MODEL to generate a protein model for each allele [84]. OmpU structures were visualized using UCSF ChimeraX [39].

### Survival assays

Overnight cultures were subcultured (1:100) in LB and grown to an optical density at 600nm (OD600nm) of 0.5. Cells were washed twice and resuspended in LB. Cultures were resuspended in LB or LB supplemented with 0.4% whole bile (Chem-Impex), 450 U/mL polymyxin B (Sigma), or 7.5μg/mL rifamycin SV (Sigma). Cultures were incubated for 1h at 37 °C in a rotary shaker after which the colony forming units were calculated by plating serial dilutions on LB agar. For exposure to RSV, cultures were incubated for 30mins. Organic acid tolerance. Exposure to organic acid was dome as previously described [41]. To investigate survival in the presence of the antimicrobial peptide P2, experiments were performed using protocol from reference 33 with minor modifications [33]. Briefly, overnight cultures of *V. cholerae* were subcultured (1:100) in LB and grown to an optical density at 600nm (OD600nm) of 0.5. Cells were washed twice with phosphate-buffered saline (PBS) pH 7.4 and resuspended in PBS pH 7.4. Aliquots of 100μL of culture were added to 400μL of PBS or PBS supplemented with P2 to achieve a final concentration of 200μg of P2. Cultures were incubated for 1.5h at 37 °C in a rotary shaker after which the colony forming units were calculated by plating serial dilutions on LB agar. The percentage of bacterial survival was calculated by comparing the c.f.u./mL of the treated versus the untreated *(N≥*3). Titrations of oxytetracycline and cinoxacin were performed using concentrations ranging from.06 – 0.5μg/mL and 7.5-15μg/mL for rifamycin SV. Cultures were incubated for either 30 min (RSV and cinoxacin) or 1h (oxytetracycline) at 37 °C in a rotary shaker after which the colony forming units were calculated by plating serial dilutions on LB agar. The percentage of bacterial survival was calculated by comparing the c.f.u./mL of the treated versus the untreated (*N*≥3).

### Phenotypic microarrays

Biolog Phenotype MicroArrays for antimicrobial chemical sensitivity, pH and osmolarity (PM 9-20) were performed in accordance with the protocol from Mackie et al. [85]. Briefly, *V. cholerae* was suspended in 1x Biolog IF-0a media with 85% transmittance (Biolog turbidimeter, 20-mm diameter tube). Following, the cells were diluted 1:200 in IF-10 media plus Biolog Redox Dye Mix D to create the inoculum (Biolog, Hayward, CA). PM 9-20 were inoculated with 100 μl/well and incubated at 37 °C with shaking. OD at 595 nm (OD_595_) was read every hour for 24h in a TECAN Sunrise microplate reader (TECAN US). Data was collected and analyzed by Magellan software (TECAN US). Biolog assays were performed in duplicate.

## ACKNOWLEDGMENTS

This work was supported by a National Science Foundation (NSF) CAREER award (#2045671) and a Burroughs Wellcome Investigator in the Pathogenesis of Infectious Disease (#1021977) to SAM.

## AUTHOR CONTRIBUTIONS

SAM designed research; TAG and MLP performed research; TAG, MLP and SAM analyzed data; TAG and SAM wrote the manuscript.

## SUPPORTING INFORMATION

**S1 Fig. Multiple sequence alignment of *ompU* alleles**. Gray boxes represent missing residues and black dots represent conserved residues. Highlighted in black boxes are identified regions of interest numbered 1-7. Green and purple dots represent alleles from Cluster 1 and Cluster 2 respectively.

**S2 Fig. OmpU Immunoblots of OmpU from representative clades**. 200ng of whole cell lysates from strains encoding alleles from (A) Cluster 1 and (B) Cluster 2 were probed using α-OmpU antibodies.

**S3 Fig. Sequence alignment and domain identification of *ompU* alleles exhibiting convergent phenotypes**. Gray boxes represent missing residues and black dots represent conserved residues. Highlighted in black boxes are confirmed regions of interest lettered A-D. Green and purple dots represent alleles from Cluster 1 and Cluster 2, respectively.

**S4 Fig. Titrations of Rifamycin SV**. Survival of *V. cholerae* N16961 WT and Δ*ompU* in the presence of varying concentrations of rifamycin SV. n≥3, \**p*<0.05.

